# Active self-touch restores bodily self-awareness following disruption by “rubber hand illusion”

**DOI:** 10.1101/2023.03.06.531301

**Authors:** Damiano Crivelli, Antonio Cataldo, Gabriella Bottini, Hiroaki Gomi, Patrick Haggard

## Abstract

Bodily self-awareness relies on a constant integration of visual, tactile, proprioceptive, and motor signals. In the “Rubber Hand Illusion” (RHI), conflicting visuo-tactile stimuli lead to changes in self-awareness. It remains unclear whether other, somatic signals could compensate for the alterations in self-awareness caused by visual information about the body. Here, we used the RHI in combination with robot-mediated self-touch to systematically investigate the role of tactile, proprioceptive, and motor signals in protecting and restoring bodily self-awareness. Participants moved the handle of a leader robot with their right hand and simultaneously received corresponding tactile feedback on their left hand from a follower robot. This self-touch stimulation was performed either before or after the induction of a classical RHI. Across three experiments, active self-touch delivered after – but not before – the RHI, significantly reduced the proprioceptive drift caused by RHI, supporting a restorative role of active self-touch on bodily self-awareness. The effect was not present during involuntary self-touch, where the participants’ hand was passively moved, suggesting that the restorative effect depends on the presence of a voluntary motor command, and that synchrony of bilateral sensory events is insufficient. Unimodal control conditions confirmed that the coordination of both tactile and motor components of self-touch was necessary to restore bodily self-awareness. These results suggest that voluntary self-touch can restore an intrinsic representation of the body following visual capture during RHI.

## Introduction

James’ description of the “same old body, always there” (1), highlights that our own body is the most familiar object in our mental life. However, it remains unclear how individual sensory experiences related to the body give rise to a general sense of body awareness, and which types of sensory signal dominate this process. An influential experimental approach to this question involves the “Rubber Hand Illusion” (RHI) (2). During the RHI, the participant receives tactile stimulation on her unseen hand, while seeing the same stimulation performed on a fake, rubber hand. If the visual and tactile stimulations are synchronous, participants often report the feeling that the rubber hand is theirs, and part of their own body: the fake hand is “embodied”. Asynchronous stimulation does not induce RHI and is commonly used as a control condition. The strength of the illusion can be measured qualitatively through questionnaires (3). Some studies have used a quantitative proxy measure by asking participants to report the position of their unseen hand (4). Participants tend to perceive their hand as shifted towards the location where they saw the fake hand. Crucially, this tendency is stronger in the synchronous than the asynchronous condition. This “proprioceptive drift” measure quantifies the visual capture of proprioception in RHI.

The RHI demonstrates that the awareness of one’s own body depends on integration of multiple sensory input signals and shows remarkable plasticity when these signals change. Further, the RHI paradigm has provided an important method for experimental studies of body awareness more generally (3). The RHI seems to show that the visual representation of body parts is more important for body awareness than truly somatic sensory inputs. Visual experience of one’s body is important for body part recognition and self-identification (5), but does not capture other key features of body awareness, namely that one’s body is sentient, and that the somatic sensations it houses are intrinsically linked to one’s sense of self.

In contrast, non-visual aspects of body awareness such as sentience and self-specificity are strongly present in the act of self-touch. Self-touch is common in humans and many other animals. It occurs in many contexts, such as grooming behaviours, affective self-stimulation, and homeostatic thermoregulation. Self-touch also occurs incidentally in the process of many other behaviours, notably feeding and bimanual object handling. A long tradition of phenomenological interest in self-touch (6, 7) focusses on the integration of motor and tactile signals in self-touch (8–10) and speculates that the resulting sensorimotor contingencies may underpin the development of a coherent, stable sense of a bodily self. This tradition emphasizes an intrinsic, or somatic sentient basis for body awareness, in contrast with the external perspective on one’s own body that lies at the heart of the RHI. The relative importance of these two components in everyday body awareness outside the laboratory remains unclear.

Some studies have combined self-touch and RHI approaches. First, Ehrsson et al. (11) used a modified form of self-touch to develop a proprioceptive analogue of the visual RHI (i.e., the somatic RHI). Participants stroked an unseen rubber hand, while receiving synchronous strokes on their own hand. This produced a convincing illusion of stroking one’s own hand, and a proprioceptive drift in the perceived position of one’s stroked hand towards the rubber hand. This result suggests that tactile and motor signals associated with self-touch make an important contribution to body awareness. This initial result was subsequently confirmed and extended (12–16). However, these studies have no visual component – participants are blindfold. It therefore remains unclear how the external visual perspective on one’s own body (as in the RHI), and the intrinsic motor-somatosensory contribution (as in self-touch) are combined.

We therefore investigated whether self-touch can restore a disturbance in bodily awareness caused by RHI. This hypothesis is motivated by findings from clinical neuropsychology. Disturbance of body awareness (17) may occur in a number of neurological and psychiatric conditions. In a single-case study of somatoparaphrenia - a rare clinical condition in which body parts are mis-attributed to others (18) - Van Stralen et al. (19) showed that actively self-touching the affected limb led to a remission of misattribution. Further, patients with hemianesthesia showed a positive modulation of tactile detection, localization, and perceived intensity when the stimuli were delivered through self- compared to other forms of tactile stimulation (20, 21).

Therefore, we have combined these two key sources of information contributing to body awareness and investigated their interaction. We first induced an alteration of body awareness using the RHI, and then investigated whether we could mitigate the effects of this intervention, and restore a normal sense of the body through self-touch. We hypothesized that active self-touch immediately after inducing an RHI would reduce the strength of the rubber hand illusion, and act to restore a ‘normal’, pre-existing awareness of the body, as previously shown in studies of neurological patients.

We used two robotic arms in a leader-follower configuration to create an artificial self-touch condition (22, 23) to test the influence of self-touch on body awareness. By moving the handle of the leader robot with their right hand, participants were able to simultaneously feel a corresponding tactile feedback on their left forearm (follower robot). Thanks to this mediated self-touch setup, and contrary to everyday self-touch involving direct skin-skin contact, the right-hand’s movement did not provide any direct spatial information about the left-hand’s position. We could therefore use the perceived position of the left hand, and particularly the well-established proprioceptive drift measure, to investigate the effects on body awareness of the RHI, of self-touch, and of the two in combination.

We report a systematic series of three experiments based on power calculation, replication, and preregistration (see Methods), to investigate the role of self-touch in bodily awareness. In Experiment 1, we explored the hypothesis that self-touch has a restorative effect on an altered body awareness caused by immediately-preceding RHI. In Experiment 2, we instead investigated if self-touch has a protective effect on body self-awareness by asking the participant to perform a self-touch stimulation immediately *before* inducing the RHI. Importantly, in both experiments we also implemented passive self-touch conditions, in which the right hand of the participant was passively moved by the experimenter while stroking the left forearm (22, 23). By comparing effects of active and passive self-touch, we could therefore investigate whether motor signals play a distinctive role in body awareness. Finally, Experiment 3 used a larger sample estimated a priori by power analysis of results of Experiment 1 and included additional unimanual controls to investigate the separate contributions of movement and touch alone to the combined effect of self-touch on body awareness.

## Results

### Does self-touch have a restorative effect on bodily self-awareness?

In Experiment 1, we tested whether brief self-touch stimulation immediately *after* the induction of the RHI could mitigate the effects of a previous RHI on body awareness. Participants (n = 16, based on a power analysis on a pilot experiment, see Methods) made a baseline proprioceptive judgement at the beginning of each trial by closing their eyes and pointing with their right hand to the location of their left hand (Figure 1A and Methods). Next, they received one of three visuo-tactile stimulation conditions: synchronous, asynchronous, and no stimulation (Figure 1B and Methods). The RHI induction phase lasted for 60 s and was followed by one of three self-touch stimulations: active self-touch, passive self-touch, and no self-touch. The self-touch stimulation was performed using coupled robots, that implemented a mediated form of self-touch, lasting 15 s (see (22, 23) and Figure 1C). Immediately after the self-touch stimulation, the participants made a second proprioceptive judgement. The participants’ proprioceptive drift (2) was computed as the difference between the final and the baseline proprioceptive judgement. Figure 2 shows the individual and average proprioceptive drift results in each condition of Experiment 1.

**Figure 1:**
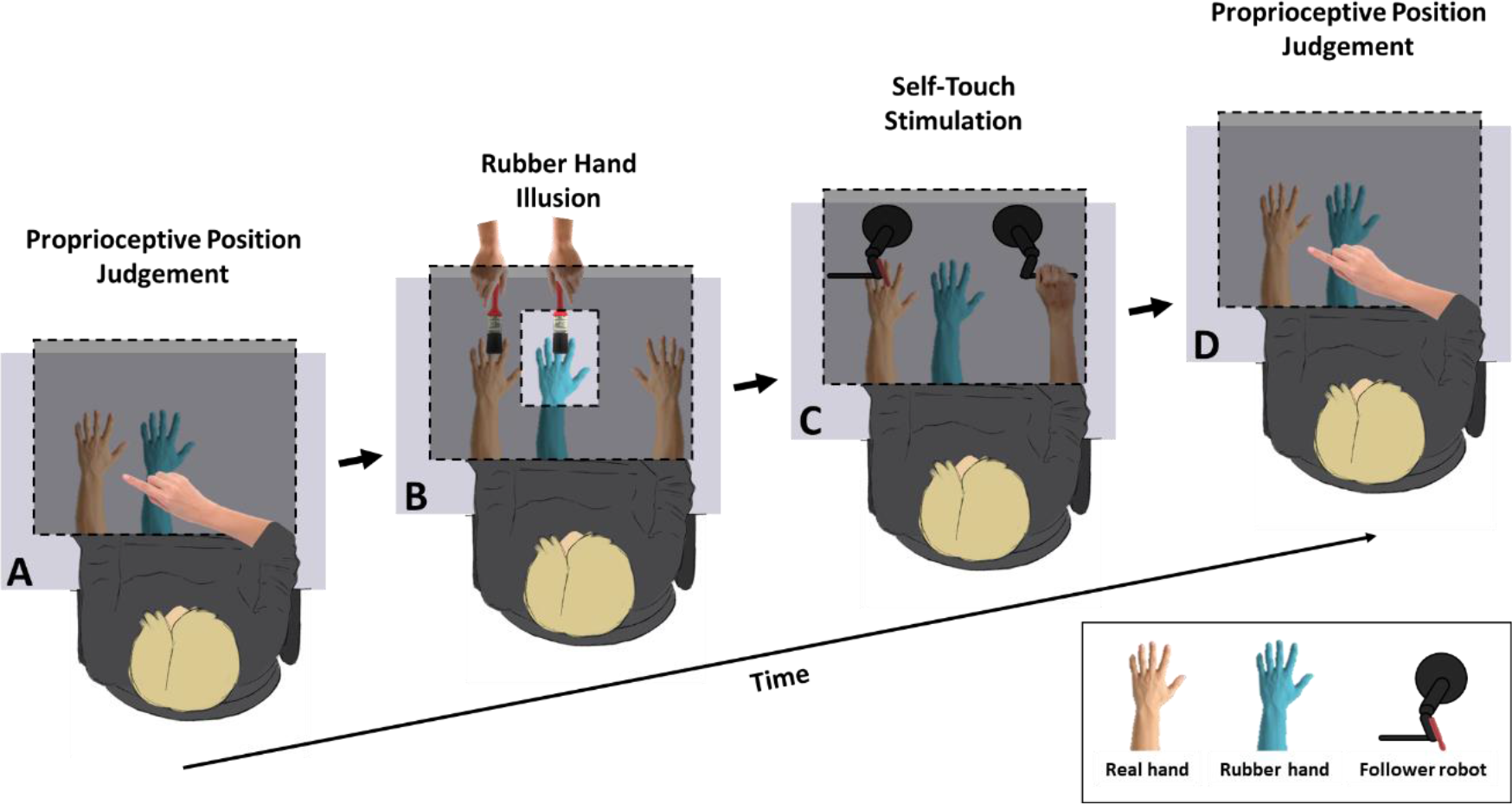
Experimental Setup. Participants sat at a desk, resting both their arms on the surface. A rubber hand was placed to the right of real left hand. The participants’ left arm and the robotic setup were covered by a foamboard screen and remained unseen throughout the entire experiment. The silicone glove was instead clearly visible through an aperture in the foamboard. **A. and D. Baseline proprioceptive judgement.** At the beginning and the end of each trial, participants were asked to close the eyes and to use their right index finger to produce a ballistic movement to point to the location immediately above the centre of their left wrist. Pointing performance was measured using a webcam suspended above the workspace. **B. RHI induction.** The experimenter sat opposite to the participant and used the two identical brushes to stroke homologous points of the participants’ left hand and the cosmetic glove either synchronously or asynchronously for one minute. **C. Self-Touch stimulation.** Self-touch was performed immediately after (Experiments 1-3) or before (Experiment 2) the RHI stimulation using two six- degrees-of-freedom robotic arms coupled in a leader-follower system. The participants were asked to move the leader robot with their right hand. The participants’ movement was reproduced by the follower robot thus generating corresponding gentle strokes from on the back of the participants’ left middle.

**Figure 2:**
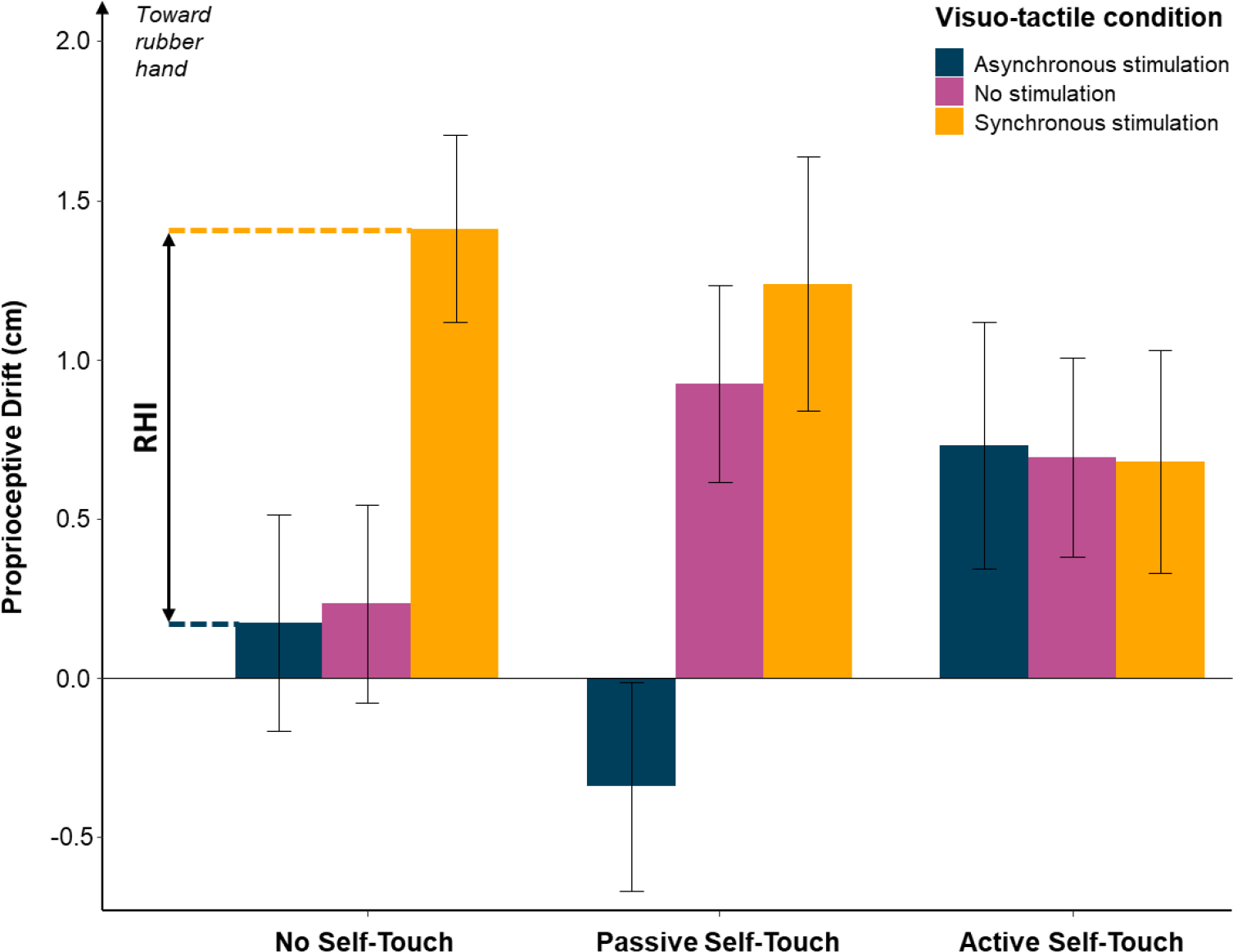
Results from Experiment 1. Both in the No Self-Touch and in the Passive Self-Touch conditions, participants showed significantly larger proprioceptive drift in the synchronous compared to the asynchronous RHI condition. In contrast, following Active Self-Touch condition, the three visuo-tactile conditions were virtually identical, suggesting a restorative effect of self-touch on bodily awareness. The error bars represent the SEM.

**Figure 3:**
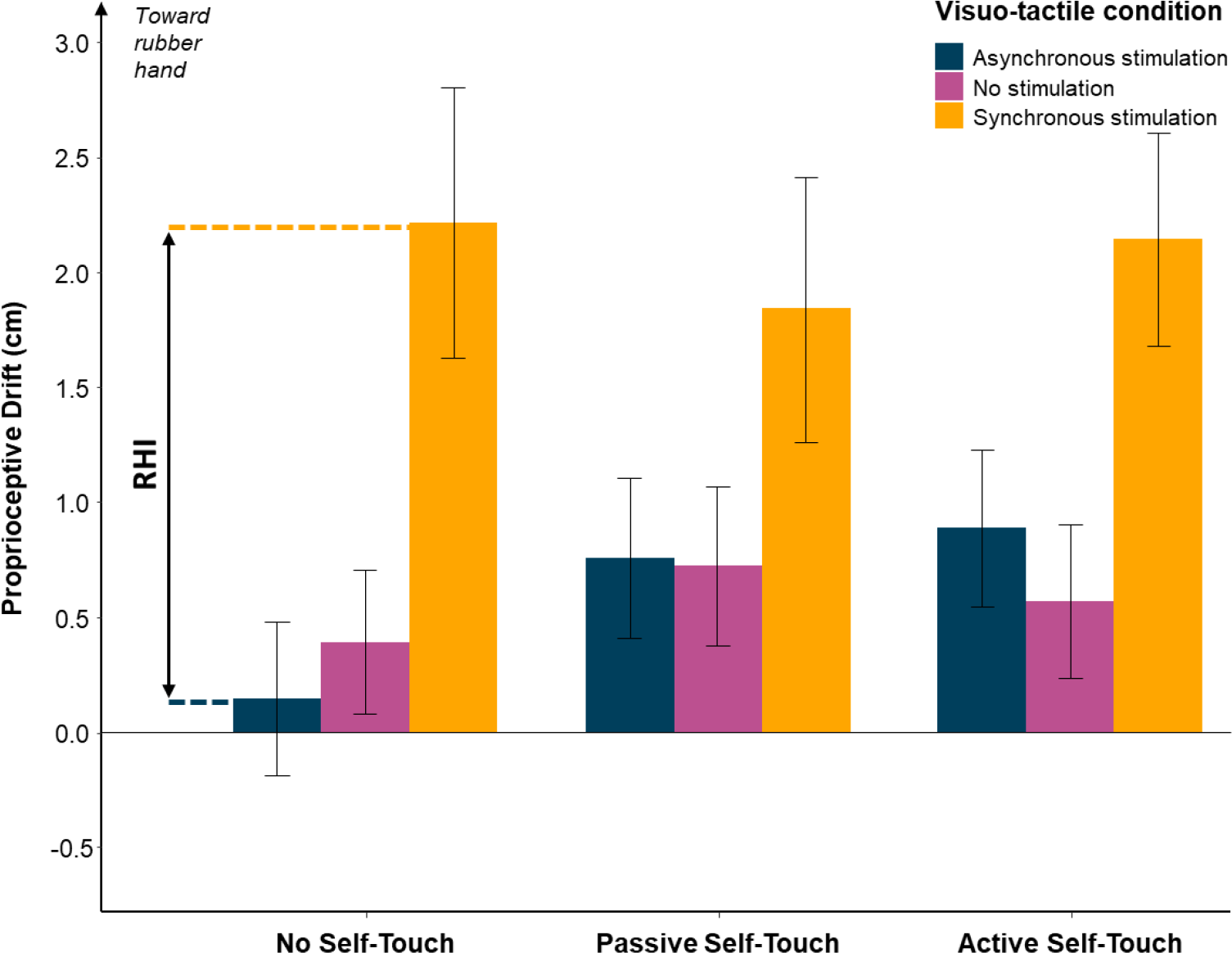
Results from Experiment 2. Self-touch delivered before the induction of the RHI had no effect on the participants’ proprioceptive drift, ruling out the hypothesis that self-touch has a protective effect on bodily awareness. The error bars represent the SEM.

**Figure 4:**
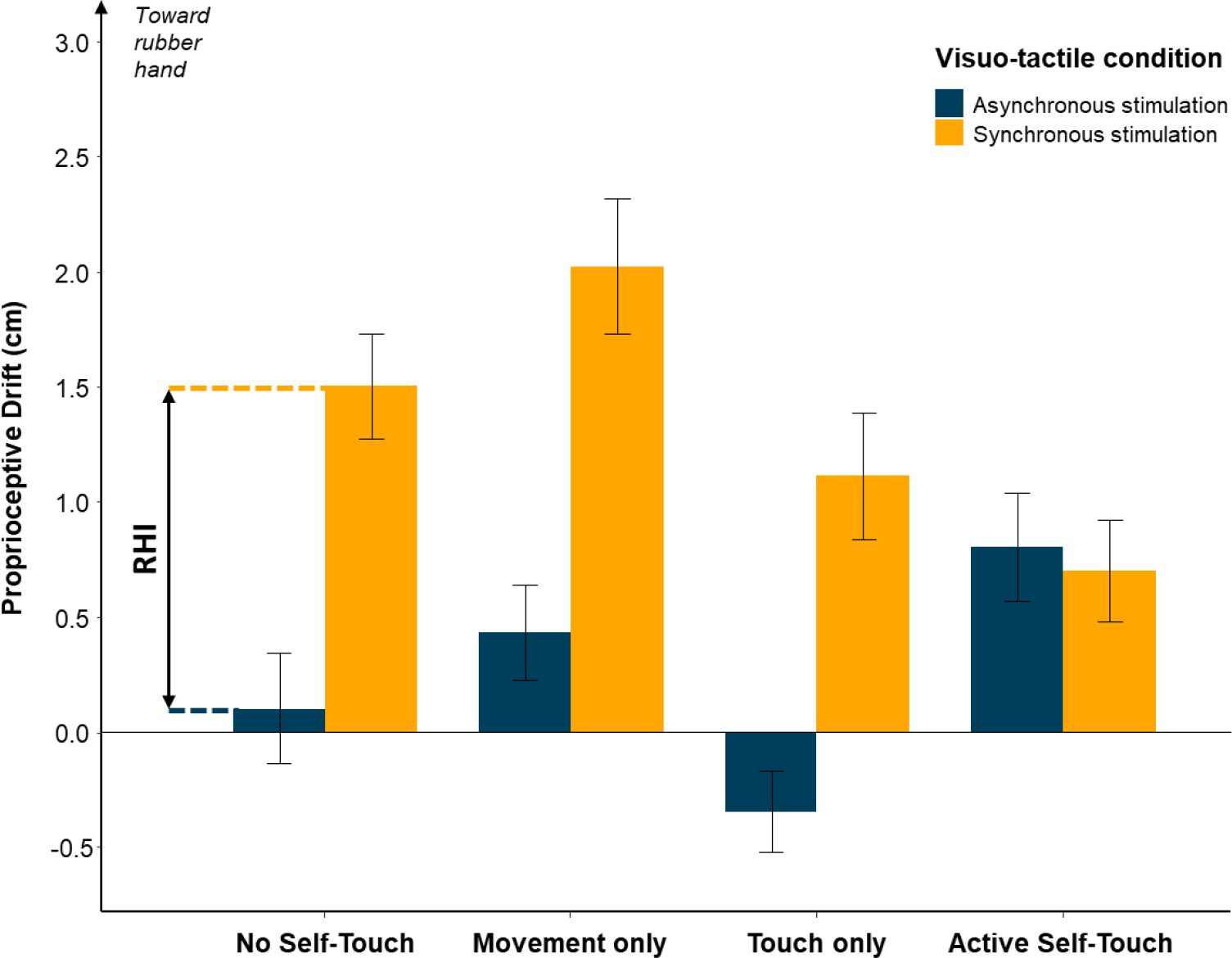
Results from Experiment 3. Motor and tactile components of self-touch were organised according to a 2 x 2 factorial arrangement, and followed immediately after synchronous or asynchronous visuo-tactile stimulation. Unimodal motor and tactile stimulation conditions and no self-touch condition all showed a significant RHI, defined as the difference in proprioceptive drive between the synchronous and asynchronous visuo-tactile conditions. After active self-touch, instead, the two visuo-tactile conditions were not statistically different, replicating the results from Experiment 1, and confirming the role of active self-touch in restoring bodily awareness. The error bars represent the SEM.

If self-touch restored body awareness, then the proprioceptive drift caused by RHI should be reduced if RHI induction is immediately followed by self-touch. In line with this hypothesis, a 3 (Visuo-tactile Stimulation: Synchronous, Asynchronous, No Stimulation) x 3 (Self-touch Condition: Active, Passive, No self-touch) repeated measures ANOVA showed a significant main effect of Visuo-tactile Stimulation induction (*F*(*2, 30*) = 11.49, *p* < .001, *η^2^p* = .434), no main effect of Self-touch Condition (*F*(*2, 30*) = 0.09, *p* = .914, *η^2^* = .006), but a marginally significant interaction between factors (*F*(*4, 60*) = 2.49, *p* = .052, *η^2^p* = .143). The trend was explained by the difference between the Synchronous and the Asynchronous in the Passive (Synchronous: *Mean* = 1.24, *SD =* 1.60; Asynchronous: *Mean* = -0.34, *SD =* 1.32; t(15) = 3.48; *p* = .003; Cohen’s d = .870) and No self-touch (Synchronous: *Mean* = 1.41, *SD =* 1.18; Asynchronous: *Mean* = 0.18, *SD =* 1.36; t(15) = 3.09; *p* = .007; Cohen’s d = .772) conditions. In contrast, the active self-touch condition was virtually identical across all the three RHI conditions (Synchronous: *Mean* = 0.68, *SD =* 1.40; Asynchronous: *Mean* = 0.73, *SD =* 1.55; No Stimulation: *Mean* = 0.69, *SD =* 1.25; *p* ≥ .929 in all cases). Thus, results from Experiment 1 provided weak evidence that active self-touch immediately *after* a RHI stimulation might mitigate the altered bodily awareness induced by the RHI, supporting a restorative role of active self-touch on bodily self-awareness.

### Does self-touch have a protective effect on bodily awareness?

In Experiment 2, we tested in a new set of participants (n = 16) whether a brief self-touch stimulation performed *before* the induction of the RHI had a *protective* effect on the participants’ bodily self-awareness, by reducing the susceptibility to the RHI. The experimental design was identical to Experiment 1, except that the order of the self-touch stimulation and the RHI induction phases was inverted. A protective effect on bodily self-awareness would be shown by a significant reduction of the participants’ proprioceptive drift after active or passive self-touch, but not after the control no self-touch condition. Contrary to this hypothesis, a 3 (Visuo-tactile Stimulation: Synchronous, Asynchronous, No Stimulation) x 3 (Self-touch Condition: Active, Passive, No self-touch) repeated measures ANOVA showed a significant main effect of Visuo-tactile Stimulation (*F(1.37, 20.57)* = 9.09, *p* = .004, *η^2^* = .377), but no main effect of Self-touch Condition (*F* = 0.69, *p* = .510, *η^2^* = .044) nor interaction between factors (*F(4, 60)* = 1.14, *p* = .347, *η^2^* = .071). The main effect of Visuo-tactile Stimulation was explained by a significantly higher proprioceptive drift in the synchronous RHI condition (*Mean* = 2.06, *SD =* 2.14) compared to both the Asynchronous (*Mean* = 0.59, *SD =* 1.38; *t(15)* = 3.32, *p* = .005; Cohen’s d = .830) and the No Stimulation condition (*Mean* = 0.56, *SD =* 1.31; *t(15)* = 3.10, *p* = .007; Cohen’s = .775), as expected.

Thus, we found no evidence that performing a brief self-touch stimulation has a protective effect against changes in body awareness induced by a subsequent RHI.

### Do unimodal components of self-touch independently affect bodily self-awareness?

Experiment 3 aimed to replicate and further investigate the results of Experiment 1. The sample size was estimated by a priori power calculation based on effects in Experiment 1, and the experiment was preregistered (see preregistration: https://osf.io/ygqnf). Given the interaction effect between Visuo-tactile Stimulation and Self-touch Condition in Experiment 1 (*η^2^* = .143) and a desired power of .80, a total of 28 participants were tested (24). Further conditions were included to investigate *why* active self-touch might restore altered bodily awareness: was restoration driven by active movement of the right hand, by tactile stimulation of the left hand, or did it require the combination of both factors? As in Experiment 1, RHI induction was always followed by a self-touch intervention. However, self-touch conditions were provided by a factorial combination of right-handovement (Present, Absent) and left-arm Touch (Present, Absent). The conditions where movement and touch were both present or both absent were identical to the active self-touch and no self-touch conditions of Experiment 1, respectively. The other two factorial combinations (movement only and touch only) served as control conditions to investigate whether either the motor or the tactile component of self-touch alone was sufficient to influence bodily awareness.

We predicted (https://osf.io/ygqnf) a significant three-way interaction between the Visuo-tactile Stimulation factor (Synchronous, Asynchronous), the Movement factor (Present, Absent), and the Touch factor (Present, Absent). This prediction was based on the hypothesis that bodily awareness, as indicated by a reduction in the proprioceptive drift associated with RHI, would be restored only by active self-touch. In particular, we hypothesized that the magnitude of the RHI- induced proprioceptive drift would be smaller when followed by an active self-touch condition involving both movement and touch, relative to other conditions. A 2 (Visuo-tactile Stimulation: Synchronous, Asynchronous) x 2 (Movement: Present, Absent) x 2 (Touch: present, absent) repeated measures ANOVA showed main effects of Visuo-tactile Stimulation (*F(1, 27)* = 50.75, *p* < .001, *η^2^* = .653), Touch (*F* = 12.18, *p* = .002, *η^2^* = .311), and Movement *F* = 4.05, *p* = .054, *η^2^* = .130, and the predicted significant three-way interactions between the factors (*F* = 18.51, *p* < .001, *η^2^* = .407). There was no mean difference in proprioceptive drift between the Synchronous and Asynchronous RHI following active self-touch (Synchronous: *Mean* = 0.70, *SD =* 1.16; Asynchronous: *Mean* = 0.81, *SD =* 1.26; *t(27)* = 0.46; *p* = .647; Cohen’s d = .088). In contrast, the difference between proprioceptive drift in synchronous and asynchronous RHI conditions was significant in all the other conditions (Movement only: Synchronous: *Mean* = 2.02, *SD =* 1.56; Asynchronous: *Mean* = 0.43, *SD =* 1.09; *t(27)* = 5.21, *p* < .001; Cohen’s d = .985; Touch only: Synchronous: *Mean* = 1.11, *SD =* 1.45; Asynchronous: *Mean* = -0.35, *SD =* 0.93; *t(27)* = 5.00, *p* < .001; Cohen’s d = .944, No Movement No Touch: Synchronous: *Mean* = 1.50, *SD =* 1.21; Asynchronous: *Mean* = .099, *SD =* 1.276; *t(27)* = 5.59, *p* < .001; Cohen’s d = 1.057). Thus, ‘complete’ active self-touch could restore the altered body awareness induced by the RHI, but the individual unimodal components of self-touch could not. This result suggests that correlation between movement and sensory stimulation associated with active self-touch is critical for its restorative effects on body awareness. Only the unique sensorimotor integration of motor and tactile information during active self-touch can restore the altered bodily self-awareness induced by the RHI.

## Discussion

Self-touch is a unique multimodal experience in which our body is both the agent that generates sensory inputs through voluntary motor action and simultaneously is the object perceived through those sensory inputs. For this reason, self-touch has long been thought to have a special significance for mental life and to be important in linking the body to a unitary, autonomous self, distinct from other objects in the environment (25, 26).

Across three experiments, we investigated the relationship between bodily self-awareness and self-touch through the rubber hand illusion (RHI). Based on previous studies on both healthy subjects (13, 15) and neuropsychological patients (19–21), we hypothesized that active self-touch could contribute to body awareness. We tested this by first inducing a well-established multisensory alteration of body awareness, namely the rubber hand illusion (RHI), and then assessing whether self-touch could either restore body awareness after these alterations or protect against subsequent alterations. Importantly, we measured body awareness via a simple, quantitative proxy measure with a clear sensory-physiological interpretation, namely the perceived position of the stimulated hand.

As we predicted, self-touch could indeed influence body awareness. Experiments 1 and 3 showed that active self-touch after the induction of a classic RHI induce a remission of the RHI effect by reducing the proprioceptive drift caused by the illusion. This complements previous findings (11, 13) who showed that a somatosensory version of the RHI could itself induce changes in body awareness, including proprioceptive drift. Crucially, our results show that active self-touch is also able to produce effects in the opposite direction, i.e., a *restoration* of bodily awareness after previous alterations induced by the RHI. This result implies that self-touch could have an important background role in reconstituting and stabilizing the representation of one’s own body as a coherent, diachronic ensemble of sensory and motor experiences.

To our knowledge, this is the first study showing a restorative effect of self-touch on this aspect of body representation in healthy subjects. However, previous studies with neurological patients are consistent with our findings.

A single case study of a stroke patient with somatoparaphrenia (19) showed that self-touch on the affected limb induced a remission of limb “disownership”. In contrast, Jenkinson et al., (27) reported no significant effects of self-touch on the sense of arm ownership in a group of patients presenting with disturbed sense of ownership (28). These authors found instead that gentle stroking by another person, designed to stimulate ‘affective touch’ CT fibres (29, 30) did reduce disturbances of ownership. The discrepancy between these studies might arise because Jenkinson et al. supported and guided the moving limb during delivery of self-touch. This was necessary since all patients were affected by hemiplegia. However, this procedure could make their self-touch intervention similar to the “passive self-touch” condition of our Experiment 1, which had no effect on body awareness. In fact, as already shown by previous studies (13, 22, 23) the presence of a voluntary motor command may be necessary to produce the distinctive effects of self-touch on body awareness. This importance of active self-stimulation and self-discovery as the key component of self-touch was also predicted on a priori grounds in previous philosophical studies (31). Our work provides the first experimental demonstration of why this might be so.

Several studies investigated effects of self-touch in hemi-anaesthesia (20, 21, 32). These suggest that self-touch modulate cross-modal attention (33). Patients perceived tactile stimuli more reliably when these were self-delivered, compared to when they were delivered by another person. This “self-touch enhancement” (20) was explained by the movements of the unaffected hand increasing the salience of events on the affected hand, like an “attentional wand” (32). Equally, explicitly drawing attention to the location of the real hand or the rubber hand can modulate the strength of RHI in healthy participants (34).

In our study, the salience of afferent signals from left hand may be decreased by the visual-proprioceptive conflict of the RHI. Indeed, Bayesian theories of multisensory integration explain proprioceptive drifts induced by the RHI as a change in the relative weightings of visual information and of proprioceptive information about left hand position. Loss of tactile sensitivity during RHI provides independent evidence that visual-tactile conflict results in downweighing of somatosensory signals from the hand (35, 36). We suggest that subsequent active self-touch might restore a normal salience to the intrinsic somatosensory signals from the left hand. Interestingly, our self-touch was mediated by a robotic system, and did not involve movement of the right hand across the midline into the space occupied by the left hand. In contrast, previous patient studies involved the hands crossing the midline to touch in contralateral space. Thus, in our study, but not in previous studies, any change in salience of signals from the left hand or redirection of attention to the left hand through self-touch was due to the motor-tactile correlations of self-touch, rather than changes in the spatial focus of attention following the as it moves, and simply tracking hand position (37). For these reasons, our effects do not seem to be a simple artefact of spatial attention. The effects of self-touch on spatial attention have not been fully explored. When one hand moves to touch another, attentional effects may be of two kinds: an ‘attentional wand’ effect of co-locating the hands in external space, and an attentional effect of motor-tactile correlation. Our robot-mediated self-touch design distinguishes these two components for the first time, and includes the latter self-touch effect, but not the former co-location effect.

We also showed that the “restorative” effect of self-touch on body awareness was not due to movement alone, nor to touch alone, but to the unique combination between these individual signals that arises in self-touch. Ruling out ‘unimodal’ explanations of self-touch effects is important for methodological reasons. For example, we measured body awareness by asking participants to point with their right hand to the location of their left hand. The simple act of moving the right arm during self-touch could potentially influence the control of these pointing movements (38) confounding motor repetition effects with our readout of body awareness. Similarly, receiving a tactile stimulus on a body part could have automatically redirected attention toward that spatial location, which could have also influenced judgements of hand position. However, these confounds are addressed by motor and tactile unimodal conditions of Experiment 3 respectively. We found that only the multimodal condition of active self-touch, and not the motor or tactile unimodal conditions, affected our measure of perceived hand location, thus controlling for these other components of sensory and motor processing, and ruling out these artefactual explanations.

Our design also distinguishes effects of self-touch from simple spatial averaging or spatial integration of the various events occurring during the experiment. Our main measure involved judgements of the proprioceptively-perceived position of the left hand. Both the rubber hand, and the participant’s right hand that administered active self-touch lay to the right of the participant’s left hand (see Figure 1). The RHI therefore involves a rightward shift in the perceived position of the left hand, as the more reliable visual evidence for hand location dominates or captures less reliable proprioceptive information. Further, any general tendency for perceived hand position to drift spontaneously towards the midline over time irrespective of stimulation (39) is controlled for by the Asynchronous stimulation condition of the RHI. The active movements of the right hand might simply attract attention towards the right. Equally, the correlation between right hand movement and left touch might produce a form of spatial attraction or spatial binding, where both events are assumed to be collocated, as is the case with every day, unmediated self-touch. However, these effects would all produce *rightward* shifts of the perceived position of the left hand. In fact, our results clearly show that active self-touch reduces or even reverses the rightward shift in perceived position of the left hand, acting in the opposite spatial direction to both the RHI and any general spatial attraction towards the right hand. Thus, our results can only be explained by self-touch causing a strengthening of body awareness with respect to the left hand, pulling the perceived position back leftwards, rather than a spatial integration effect analogous to RHI.

Participants also answered a series of explicit questions about body awareness. Our analyses showed no effects of active self-touch on these explicit measures (see Supplemental Information). Dissociations between questionnaire responses and quantitative measures based on perceived position are common in the body awareness literature (4, 40–43). Recent accounts of somatoparaphrenia (44) consider the erroneous spatial representation of the limb position, due to a poor proprioceptive update, as the key component of the deficit. Measures of perceived location have the advantage of being implicit, and do not rely on participants’ comprehension of the words used in body awareness questionnaires (3, 45). Further, theoretical considerations suggest that spatial localization is central to constructing a mental representation of one’s own body (46).

Our measures of body awareness were based on proprioceptively-perceived position of the left hand. Since the left hand did not move at any point during the experiment, the effects of self-touch on body awareness cannot reflect changes in any proprioceptive afferent signalling. Self-touch must therefore have changed a *central* state estimate of hand position based on proprioceptive input from the left arm. Bayesian interpretations of the RHI (47) suggest that proprioceptive drift in the RHI can be reduced or reversed by increasing the precision of proprioceptive signals of hand position (or decreasing the precision of the visual signal) (cf. 48). We conclude that active self-touch enhances proprioception from the touched limb, potentially by improving proprioceptive signal-to-noise ratios. The physiological mechanisms underlying this multisensory interaction require further investigation. Close integration of tactile and proprioceptive information is found in many post-primary somatosensory cortical regions, such as posterior parietal cortex (49). However, the crucial additional role of voluntary movement of the contralateral hand in our experiment’s points to a neurophysiological mechanism beyond mere tactile-proprioceptive integration (50). In general, our results emphasize the strong importance of voluntary self-touch for body awareness (13, 50–52). Therefore, central, cognitive manipulations of proprioceptive processing, such as those triggered by self-touch, may offer therapeutic promise in restoring disturbances in body awareness that occur in a range of neurological and psychiatric diseases.

### No evidence for a protective effect of self-touch on bodily self-awareness or bodily ownership

Surprisingly, Experiment 2 showed no effect of active or passive self-touch on the strength of a subsequent RHI, assessed either via proprioceptive drift, or by direct questions. In contrast, Lesur et al. (15) previously showed that self-touch could prospectively protect the sense of body ownership from mismatching signal. Some key differences between our study and theirs could however account for this discrepancy. First, their study used temporal mismatch of the visuo-motor signals, which are known to induce feelings of dis-embodiment through self-touch itself. Moreover, their VR setup allowed participants to see their hand throughout, whereas our participant’s left hand was always hidden.

## Limitations

One limitation of the present study is that we did not provide the skin-to-skin self-touch of everyday experience, but instead, a robot-mediated analogue of self-touch (22, 23). This modification was necessary to ensure that the position of the left hand (the key readout for our proprioceptive drift measures of bodily awareness) could not be trivially known from the proprioceptive signals arising from the right arm. Further, mediated self-touch is common during everyday life (e.g., hairbrushes) and does not destroy the core experiences of self-stimulation, affective self-affirmation, and coherence of bodily self.

## Materials and Methods

### Participants

The final sample size for Experiments 1-2 (n = 16) was decided a priori based on a power analysis on the results of a pilot study (n = 8) with a similar design. In the pilot study, the effect size for the interaction of Self-touch Condition (active, passive) and Visuo-tactile Stimulation (Synchronous, Asynchronous) was dz = -.848, considered to be very large using Cohen’s criteria (53). With an alpha = .05 and power = .8, the projected sample size indicated to demonstrate a main effect of type of movement was 13 participants (24). We nevertheless set a sample size of n = 16 to ensure sufficient statistical power.

The final sample size for Experiment 3 (n = 28) was determined through a power analysis on the results of Experiment 1, where the effect size for the interaction of Self-touch Condition and Visuo-tactile Stimulation was η^2^p = .143. With an alpha = .05 and power = .8, the projected sample size indicated to demonstrate an interaction effect of type of movement and RHI was 28 participants (24).

A total of 60 healthy volunteers (45 females; age between 18 and 31) were recruited for the study. All participants were right-handed, had normal or corrected-to-normal vision, and had no previous history of mental or neurological illness. The experimental protocol was approved by the Research Ethics Committee of University College London and adhered to the ethical standards of the Declaration of Helsinki. All participants were naïve regarding the hypotheses underlying the experiment and provided their written informed consent before the beginning of the testing, after receiving written and verbal explanations of the purpose of the study. All participants received monetary compensation (£8 per hour) for their involvement in the study. The hypotheses, procedures, and analyses of Experiment 3 were pre-registered (https://osf.io/ygqnf).

### Apparatus

Figure 1 shows a schematic representation of the experimental setup. Participants sat at a desk, resting both their arms on the surface. A left cosmetic silicone glove (Realistic Prosthetics Ltd., model RPL 503/505, UK) filled with cotton wool was placed in front of the participants at ∼20 cm(54) to the right of their left hand so to be aligned to their body midline. The participants’ left arm and the robotic setup were covered by a foamboard screen and remained unseen throughout the entire experiment. The silicone glove was instead clearly visible through an aperture in the foamboard. A desk lamp was used to illuminate the rubber hand under the foamboard.

The RHI was elicited using two identical brushes, following the classical RHI procedure (2, 55). The experimenter sat opposite to the participant and used the two brushes to stroke homologous points of the participants’ left hand and the cosmetic glove (between the middle and the index finger) either synchronously (∼1 Hz) or asynchronously (∼1 Hz, 180° out of phase). The RHI stimulation lasted for 60 s (55). To obtain an estimate of the participants’ proprioceptive drift, a webcam was mounted on the ceiling above the setup. The webcam provided a top view of the participants’ arms and was used to take accurate measurements of the pointing movements made by the participants at the end of the RHI induction (see Figure 1). The coordinates of each proprioceptive judgement were extracted by each picture through the ImageJ software (http://rsbweb.nih.gov/ij/) and then converted from pixels to centimetres.

The sensorimotor self-touch stimulation was implemented using two six-degrees-of-freedom robotic arms (3D Systems, Geomagic Touch X, South Carolina, USA) linked in a computer-controlled leader-follower system (see Figure 1). In this system, any 3D-movement of the right-hand leader robot is reproduced by the follower robot with an estimated lag of ∼2.5 ms (22, 23). A wooden rod with a rounded tip was attached to the follower robot in correspondence of the middle finger of the participants’ left hand. Thus, the leader robot movements produced corresponding gentle strokes from the follower robot on the back of the participants’ left middle finger. This setup allowed us to create a laboratory-equivalent of ordinary tool-mediated self-touch. Importantly, the use of tool-mediated self-touch as opposed to skin-to-skin self-touch allowed us to make sure that the right-hand movement did not provide any spatial information about the position of the left-hand.

### Experimental design

Experiments 1 and 2 tested, respectively, whether self-touch has a restorative vs. a protective effect on bodily self-awareness. We reasoned that if self-touch has a restorative effect on bodily self-awareness, a reduction of RHI effect should be observed when a brief self-touch stimulation is performed *after* the RHI induction (Experiment 1). Conversely, if self-touch has a protective effect on bodily self-awareness, a brief self-touch stimulation should reduce the participants’ susceptibility to a subsequent RHI (Experiment 2). Both experiments had a 3 (Visuo-tactile Stimulation: Synchronous, Asynchronous, No Stimulation) x 3 (Self-touch Condition: Active, Passive, No self-touch) within-participants design. The order of both factors was completely randomized across participants. Each of the nine possible combinations of these factors was repeated three times, giving a total of 27 trials per participant for a testing session of about ∼90 min. Between trials, the participants were asked to take short breaks and to make large movements with their arms so to cancel out any lingering effect from the previous trial.

Experiment 3 aimed to replicate the findings of Experiments 1 in a larger sample and to provide further experimental control conditions for the potential unimodal effect of right-hand movement and left-hand touch alone. The experiment had a 2 (Visuo-tactile Stimulation: Synchronous, Asynchronous) x 2 (Movement: Present, Absent) x 2 (Touch: present, absent) fully within-participants factorial design. Each of the eight combinations of these factors was repeated three times, for a total of 24 trials and a testing session of about ∼90 min. Short breaks were introduced between trials to prevent any potential lingering effect from the previous stimulation.

### Procedure

Each trial in each experiment consisted of a RHI induction phase and a self-touch stimulation phase which was performed either after (Experiments 1 and 3) or before (Experiment 2) or the RHI (see Figure 1). Additionally, participants performed two pointing movements, one at the beginning of the trial (baseline pointing), and one at the end of the trial (late pointing). Before the beginning of each experiment, participants familiarized with the experimental setup and received specific training for each phase of the experiment.

#### Experiment 1

At the beginning of each trial, participants performed a baseline pointing movement. They were asked to close the eyes and to use their right index finger to produce a ballistic movement starting from the resting position on the desk and landing on the foamboard, in correspondence of the middle of their left wrist. Participants were asked to be as accurate as possible and to keep their finger on the foamboard for a few seconds to allow the experimenter to take a picture of their pointing through the webcam placed above the setup. Next, in the RHI induction phase, participants were asked to open their eyes and to fixate the cosmetic glove through the aperture in the foamboard, while the experimenter performed either a synchronous or asynchronous RHI stimulation for one minute. In a third (control) condition, participants fixated the rubber hand for one minute without receiving any visuo-tactile stimulation. Right after, in the self-touch phase, participants were asked to grasp the handle of the leader robot with the right hand and then close their eyes. In the active self-touch condition, participants produced short (∼6 cm) back-and-forth movements on the proximo-distal axis. In the passive self-touch condition, participants grasped the robot handle with their right hand and then rested as passively as possible while the experimenter moved the handle of the leader robot in the same fashion as the active self-touch condition described above. In both conditions, the follower robot generated simultaneous and spatially corresponding tactile strokes on the back of participants’ left middle finger. Either self-touch stimulation lasted for 15 s, as (55) showed a consistent presence of proprioceptive drift 15 sec after a 60 sec RHI. In a third (control) condition, no self-touch stimulation was provided, and the participants were asked to just wait for 15 s with their eyes closed before moving to the next phase of the trial.

Finally, the participants were asked to close the eyes again and to perform another pointing movement to obtain an estimate of their proprioceptive drift. At the end of the first trial of each condition, participants also filled in a short questionnaire on their subjective experience of the RHI (3) (see also Supplemental Information).

#### Experiment 2

Experiment 2 was identical to Experiment 1 in all respects, except that the order of the self-touch and the RHI phases was inverted, such that participants first performed one of the three self-touch stimulations and then experienced one of the three RHI conditions. As in Experiment 1, a baseline and a final pointing movement were acquired at the beginning and the end of each trial, providing an estimate of the participants’ proprioceptive drift. RHI questionnaires were delivered at the end of the first trial of each condition.

#### Experiment 3

In Experiment 3, the order of the self-touch and RHI phases was the same as in Experiment 1. However, the active and passive self-touch conditions were replaced by a factorial combination of Movement (Present, Absent) and Touch (Present, Absent). The conditions where movement and touch were both present or both absent were identical, respectively, to the active self-touch and no self-touch conditions in Experiment 1. The other two factorial combinations (movement only and touch only) served as control conditions to investigate whether either component of self-touch was sufficient to mediate any effect on bodily self-awareness. In the movement only condition, participants held the handle of the leader robot with their right hand and performed the same proximo-distal movements described above. Crucially, the follower robot was disconnected from the leader-follower system in this condition, such that the participants’ movement did not produce any tactile stimulation. In the touch only condition, instead, the participants were asked to rest their hand on the desk while the experimenter moved the follower robot. This produced a tactile stimulation on the participants’ left hand in absence of any right-hand movement. As in the other experiments, a baseline and a final pointing movement were acquired at the beginning and the end of each trial, providing an estimate of the participants’ proprioceptive drift. RHI questionnaires were delivered at the end of the first trial of each condition.

### Statistical analysis

We operationalized participants’ bodily self-awareness in terms of proprioceptive drift (2, 4), defined as the perceived position of the participant’s real left hand on the mediolateral axis. This measure was acquired as a series of independent proprioceptive judgements, performed through pointing movements. The proprioceptive drift was expressed as the difference between the perceived position before the start of the trial (i.e., baseline) and the one expressed after each RHI and self-touch manipulation.

To test our hypothesis that active self-touch has a restorative (Experiment 1) or protective (Experiment 2) effect on bodily self-awareness, we ran two separate 3 (Visuo-tactile Stimulation: Synchronous, Asynchronous, No Stimulation) x 3 (Self-touch Condition: Active, Passive, No self-touch) repeated measures ANOVAs on the proprioceptive drift scores and self-reports of all participants in Experiments 1 and 2. Similarly, we ran a 2 (Visuo-tactile Stimulation: Synchronous, Asynchronous) x 2 (Movement: Present, Absent) x 2 (Touch: present, absent) repeated measures ANOVA on the proprioceptive drift scores and self-reports in Experiment 3 to replicate Experiment 1 and control for the unimodal effects of movement and touch alone. ANOVAs were followed up by post-hoc pairwise comparisons when appropriate. When necessary, pairwise comparisons were Holm-Bonferroni corrected.

To assess that the assumptions of the linear model were not violated, we checked that the residuals of the models were normally distributed by visually examining Q–Q plots (see Supplemental Information).

### Data and materials availability

All data needed to evaluate the conclusions in the paper are present in the paper and/or the Supplemental Materials. Raw data associated with this manuscript can be found at in the accompanying Open Science Framework repository, at https://osf.io/ygqnf.

## Acknowledgments

This study was supported by funding from NTT (The Nippon Telegraph and Telephone Corporation, Japan) to P.H and H.G. A.C and P.H were funded by the European Union’s Horizon 2020 research and innovation programme under grant agreement No 101017746.

