## Supplemental Information for "Active self-touch restores bodily self-awareness following disruption by “rubber hand illusion”" (from another version of this preprint: version 1, 2023-03-08)

**1. Supplemental results of the embodiment questionnaires**

At the end of each trial, participants were asked to fill in a short questionnaire on their subjective experience of the RHI. Participants were asked to indicate their agreement or disagreement with the following statements by circling the corresponding number: 1) Strongly disagree, 2) Disagree, 3) Partly disagree, 4) Neither agree nor disagree, 5) Partly agree, 6) Agree, 7) Strongly agree”:

- (Q1) It felt as if the rubber hand was my hand.
- (Q2) It felt as if I was looking directly at my left hand.
- (Q3) It felt as if I might have more than one left hand or arm.
- (Q4) The rubber hand began to resemble my (real) left hand.
- (Q5) It felt as if my (real) left hand were turning ‘rubbery’.
- (Q6) It seemed as though the touch I felt was caused by the paintbrush touching the rubber hand.
- (Q7) It seemed as if I were feeling the touch of the paintbrush in the location where I saw the rubber hand touched.

To test our hypothesis that active self-touch has a restorative (Experiment 1) or protective (Experiment 2) effect on bodily self-awareness, we ran two separate 3 (Type of RHI: Synchronous, Asynchronous, No RHI) x 3 (Type of Self-touch: Active, Passive, No self-touch) x 7 (Question number: Q1, Q2, Q3, Q4, Q5, Q6, Q7) repeated measures ANOVAs on the self-reports score of all participants in Experiments 1 and 2. Similarly, we ran a 2 (Type of RHI: Synchronous, Asynchronous) x 2 (Movement: Present, Absent) x 2 (Touch: present, absent) x 7 (Question number: Q1, Q2, Q3, Q4, Q5, Q6, Q7) repeated measures ANOVA on the self-reports score in Experiment 3 to further investigate a possible restorative effect of Self-Touch on bodily self-awareness while also controlling for the unimodal effects of movement and touch alone. ANOVAs were followed up by contrast and post-hoc pairwise comparisons when appropriate. When necessary, pairwise comparisons were Holm-Bonferroni corrected.

### *Experiment 1*

Mauchly's test of sphericity indicated that the assumption of sphericity was violated so we applied Greenhouse-Geisser Correction. Experiment’s 1 (Tab 1) showed a significant main effect of Type of RHI induction (*F_(2, 176)_* = 157.709, *p* < .001, *η^2^_p_* = .6) and Question number (*F_(6, 105)_* = 3.494, *p* < .003, *η^2^_p_* = .166), but no main effect of Type of Self-touch (*F_(4, 161)_* = 2.49, *p* = .085, *η^2^_p_* = .023). The main effect of the Type of RHI factor was explained by the fact that subjects generally reported higher score in the Synchronous (*Mean* = 2.92, *SD =* ±2.17) compared to the Asynchronous *(Mean* = 1.42, *SD =* ±1.70) and No self-touch (*Mean* = 0.551, *SD =* ±1.18) conditions (see Tab 2). Moreover, the interaction between Type of RHI induction and Question number was significant (*F_(10, 176)_* = 6.639, *p* < .001, *η^2^_p_* = .275), confirming that the different statement of the embodiment questionnaire reflect different aspects of the embodiment experience (Tab 3, Fig 1). Critically, all the other interaction were not significant, revealing no restorative effect of Self-Touch on the explicit component of the RHI.

**Supplemental Table 1**: ANOVA table of Experiment 1

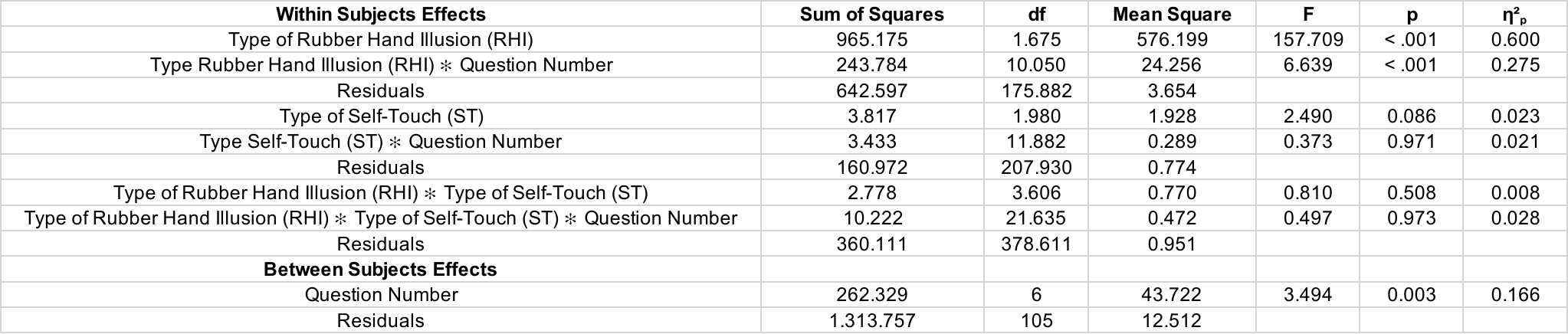

**Supplemental Table 2**: Post-Hoc comparison of Experiment 1

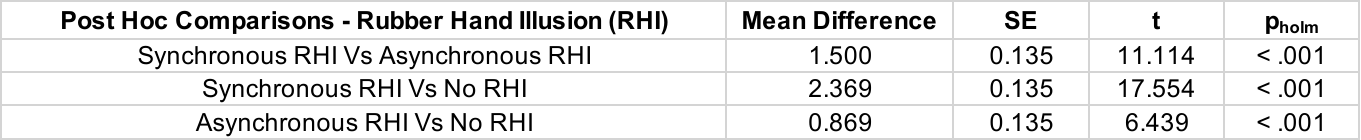

**Supplemental Table 3**: Custom contrast of Experiment 1

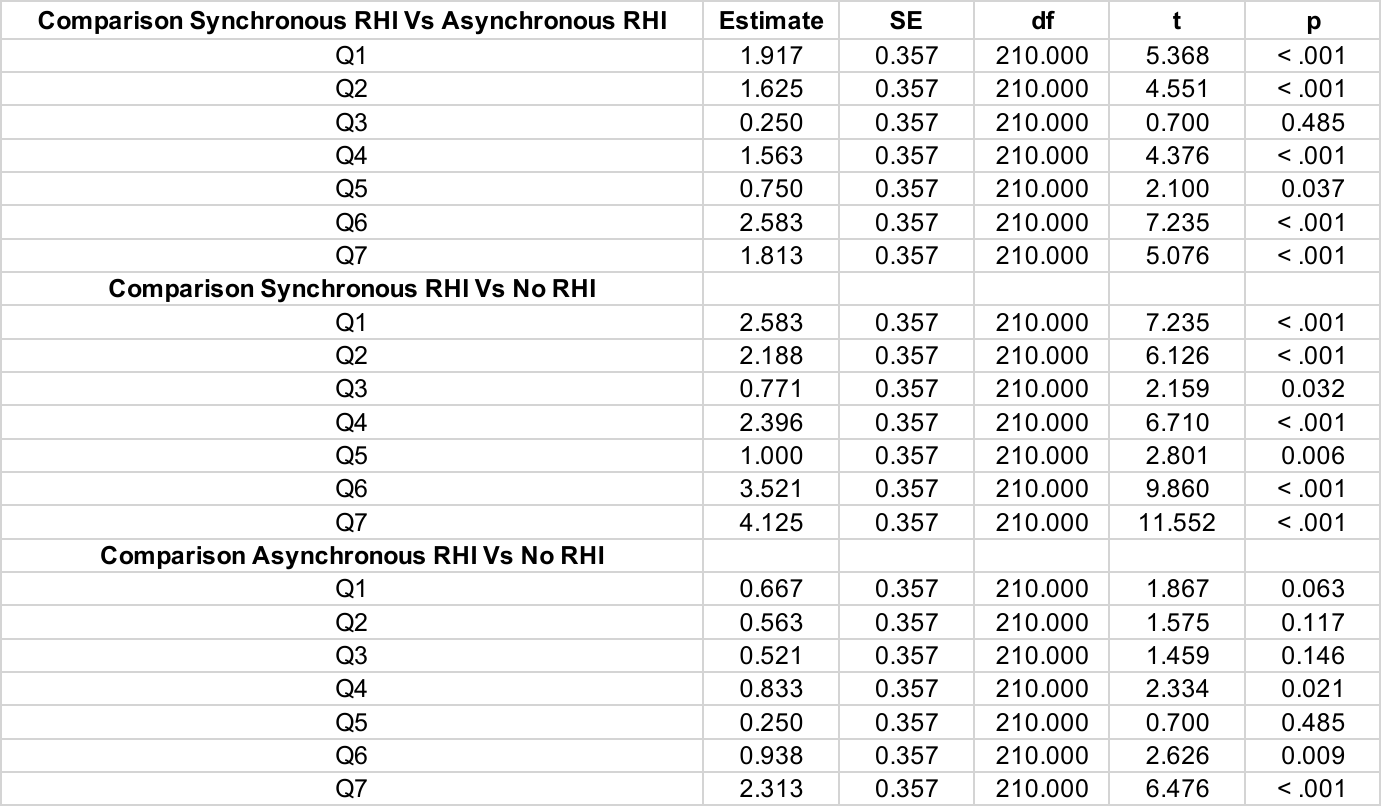

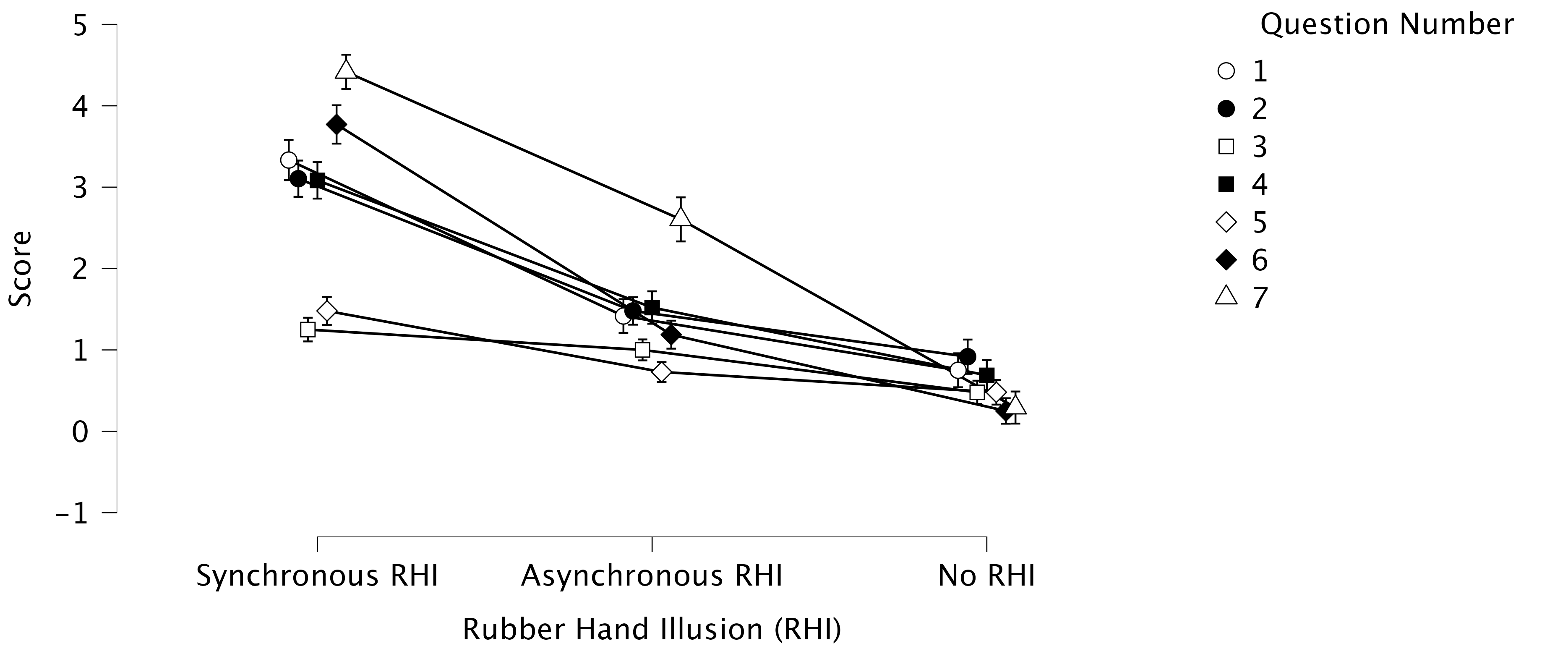

**Supplemental Figure S1**: The plot shows the mean score and the standard error of each question scores in different RHI conditions of Experiment 1.

### *Experiment 2*

Mauchly's test of sphericity indicated that the assumption of sphericity was violated so we applied Greenhouse-Geisser Correction. Experiment’s 2 (Tab 4) showed a significant main effect of Type of RHI induction (*F_(2, 205)_* = 151.617, *p* < .001, *η^2^_p_* = .591) and Type of Self-touch (*F_(2, 207)_* = 3.644, *p* = .029, *η^2^_p_* = .034), but no main effect of Question number(*F_(6, 105)_* = 1.791, *p* < .108, *η^2^_p_* = .093). The main effect of the Type of RHI factor reflect the same trend of experiment 1 *(*Synchronous RHI: *Mean* = 3.77, *SD =* ±2.10; Asynchronous RHI: *Mean* = 2.31, *SD =* ±1.98; No RHI: *Mean* = 1.34., *SD =* ±1.77; see Tab 5), while the main effect of Self-Touch was explained by the fact that subjects generally reported higher score in the Active Self-Touch condition (*Mean* = 2.56, *SD =* ±2.26) and the Passive Self-touch *(Mean* = 2.52, *SD =* ±2.14; see Tab 5) compared to the No Self-Touch *(Mean* = 2.33, *SD =* ±2.18; see Tab 5) condition. Similarly to experiment 1, the interaction between Type of RHI induction and Question number was significant (*F_(12, 206)_* = 6.782, *p* < 0.001, *η^2^_p_* = .279; Tab 5, Fig 2). Critically, all the other interaction were not significant, revealing no specific protective effect of Self-Touch on the explicit component of the RHI.

**Supplemental Table 4:** ANOVA table of Experiment 2

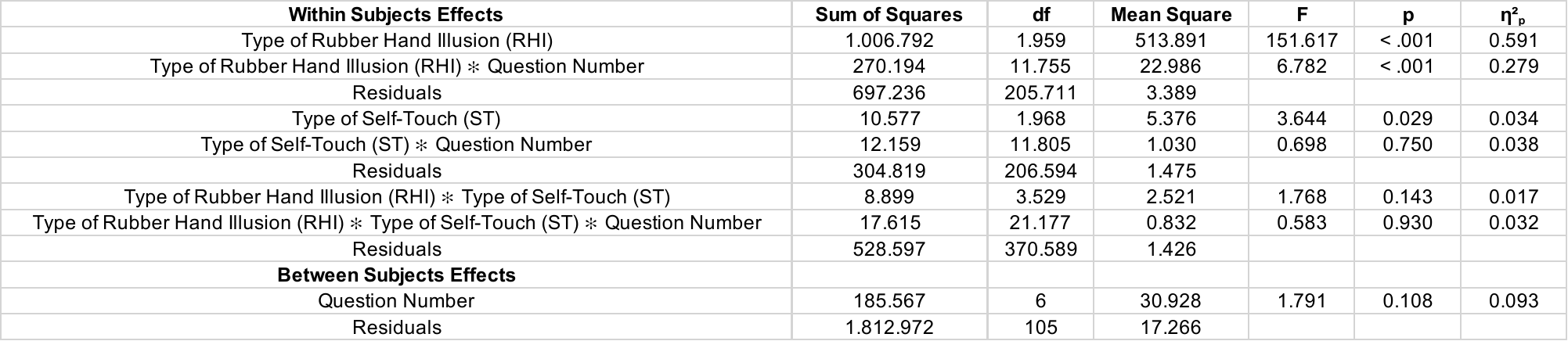

**Supplemental Table 5:** Post-Hoc comparison of Experiment 2

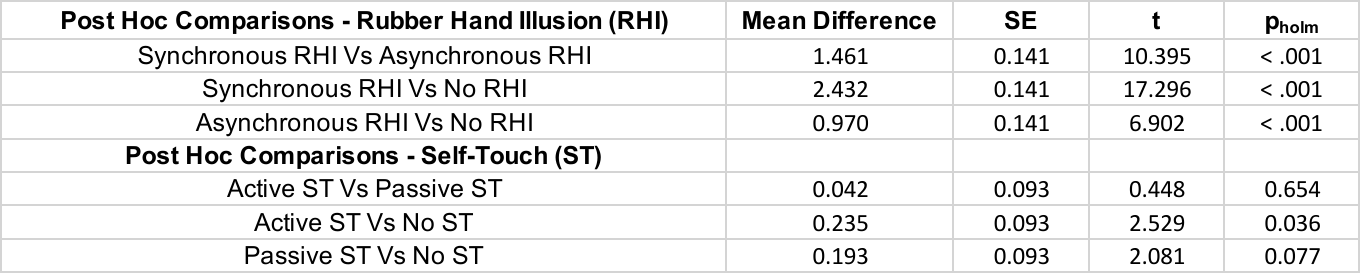

**Supplemental Table 6:** Custom contrast of Experiment 2

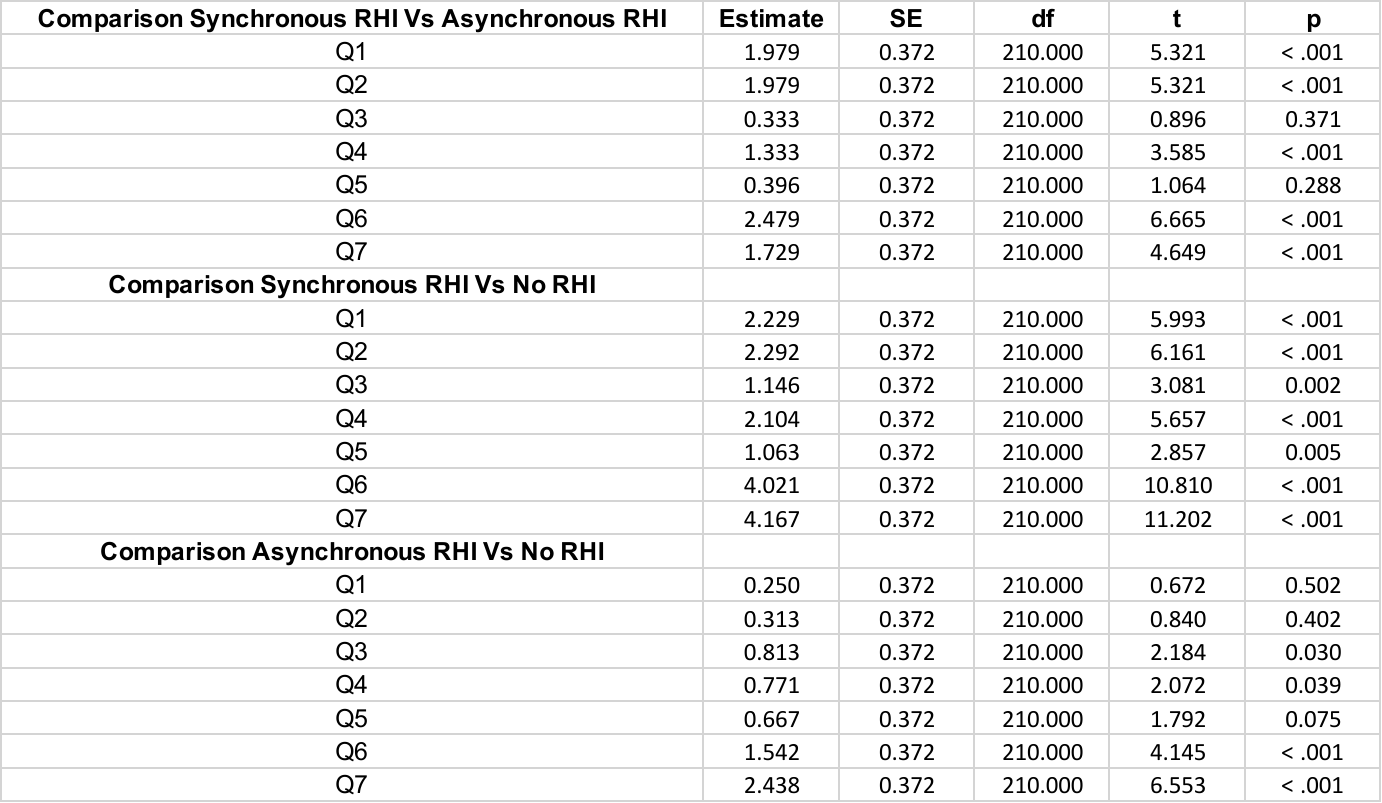

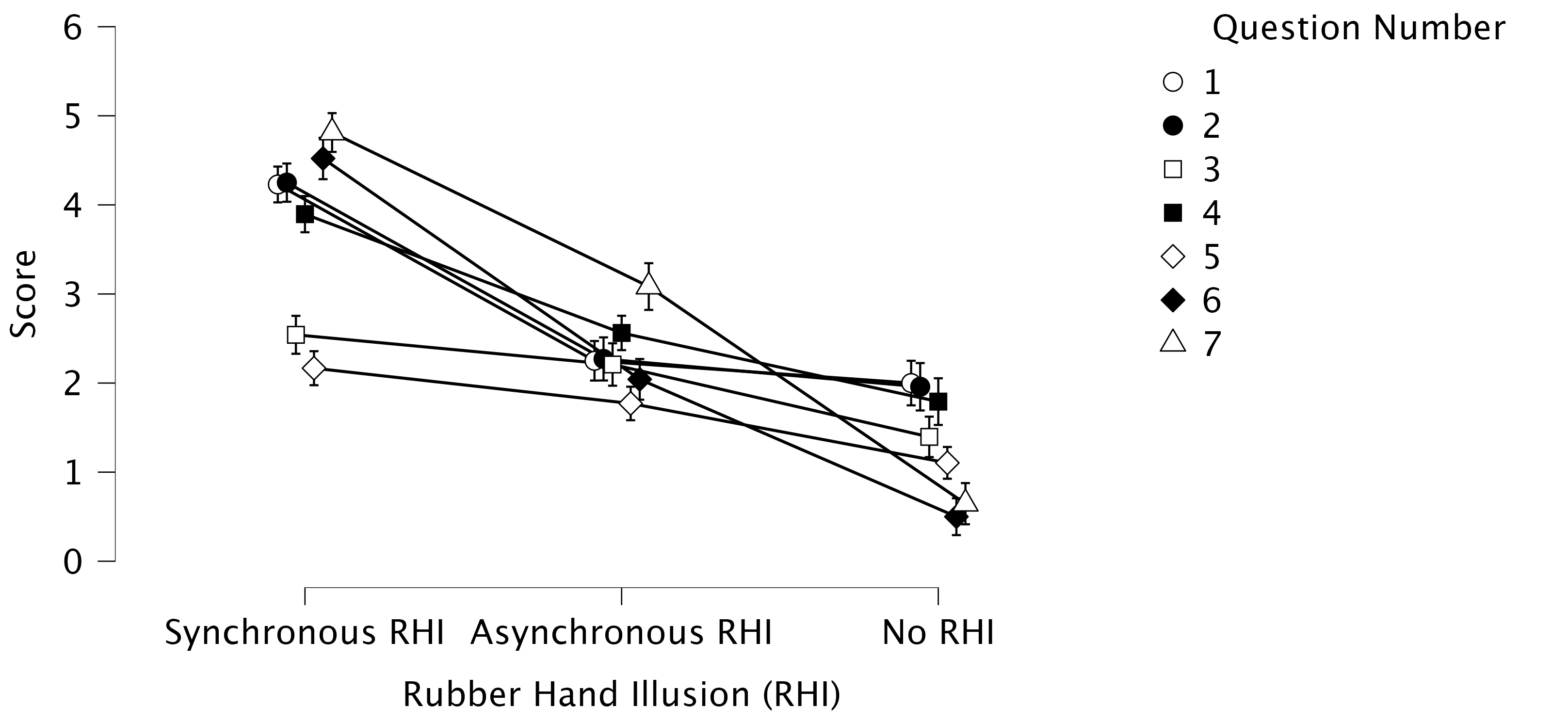

**Supplemental Figure 2:** The plot shows the mean score and the standard error of each question scores in different RHI conditions of Experiment 2.

### *Experiment 3*

Experiment’s 3 (Tab 7) showed a significant main effect of Type of RHI induction (*F_(1, 189)_* = 151.003, *p* < .001, *η^2^_p_* = .444) and Question number(*F_(6, 189)_* = 9.436, *p* < .001, *η^2^_p_* = .231), but no main effect of Touch (*F_(1, 189)_* = 1.380, *p* = .242, *η^2^_p_* = .007) or Movement (*F_(1, 189)_* = 2.796, *p* = .096, *η^2^_p_* = .015). The main effect of the Type of RHI factor was explained by the fact that subjects generally reported higher score in the Synchronous (*Mean* = 3.15, *SD =* ±2.02) compared to the Asynchronous *(Mean* = 1.99, *SD =* ±1.78; see Tab 2). Again, the interaction between Type of RHI induction and Question number was significant (*F_(6, 189)_* = 5.914, *p* < .001, *η^2^_p_* = .158) (Tab 3, Fig 1). Interestingly, results showed an interaction of both Touch (*F_(6, 189)_* = 29.736, *p* < .001, *η^2^_p_* = .136) and Movement (*F_(6, 189)_* = 20.99, *p* < .001, *η^2^_p_* = .100) with Type of RHI induction. This interaction effect revealed that the difference between the Synchronous and Asynchronous conditions was smaller when Touch (Touch present-Synchronous: *Mean* = 3.04, *SD =* ±2.07, Touch present-Asynchronous: *Mean* = 2.17, *SD =* ±1.80) or Movement (Movement present-Synchronous: *Mean* = 2.98, *SD =* ±2.04, Movement present-Asynchronous: *Mean* = 2.07, *SD =* ±1.78; ) were present compared to when they were absent (Touch absent-Synchronous: *Mean* = 3.26, *SD =* ±1.96, Touch absent-Asynchronous: *Mean* = 1.80, *SD =* ±1.74; Movement absent-Synchronous: *Mean* = 3.32, *SD =* ±1.99, Movement absent-Asynchronous: *Mean* = 1.91, *SD =* ±1.78; see Fig 4 and 5). However, the effect was significant for all questions, including questions number 3 and 5 that are usually considered “control” questions. Therefore, it is unclear whether this result reflects a specific effect of touch on explicit bodily awareness or rather a spurious response bias effect. Critically, the quadruple interaction between Type of Rubber Hand Illusion, Movement, Touch and Question number was not significant, thus suggesting no effect of Self-Touch on the explicit component of the RHI. All the other interactions were not significant.

**Supplemental Table 7:** ANOVA table of Experiment 3

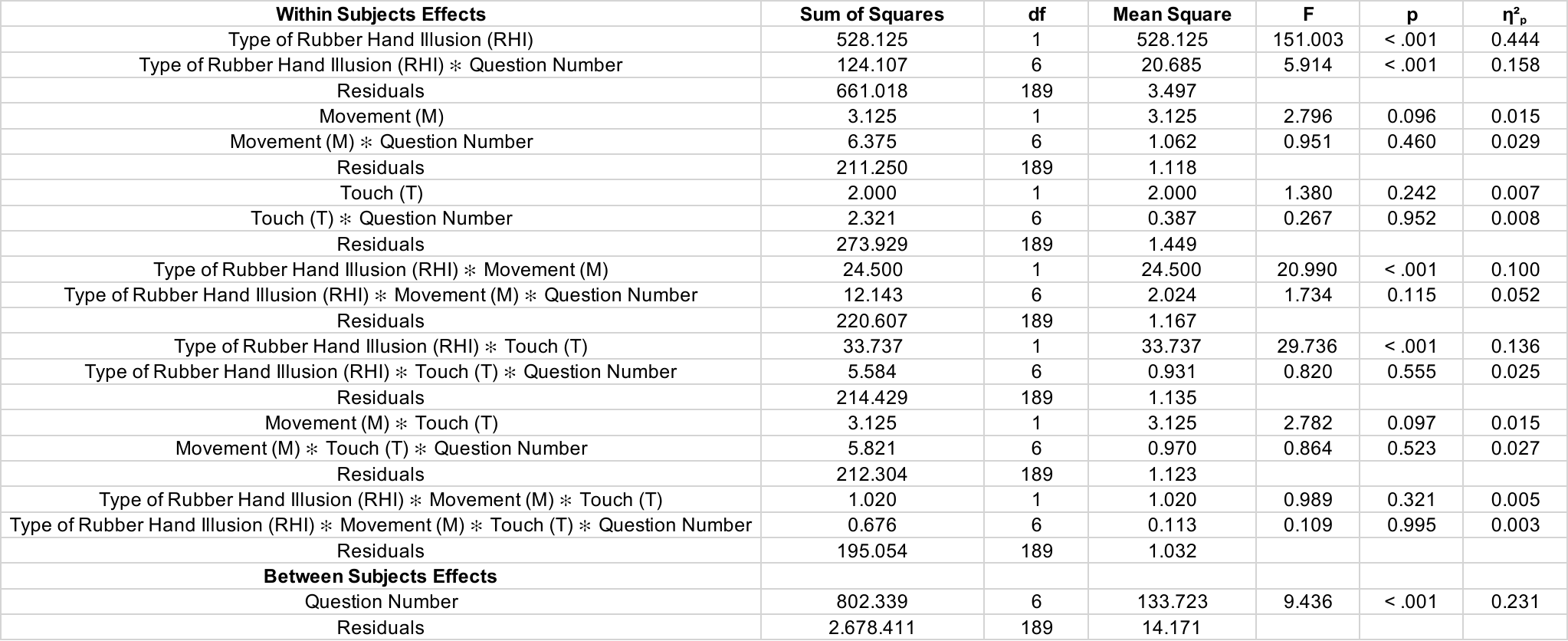

**Supplemental Table 8:** Post-Hoc comparison of Experiment 3

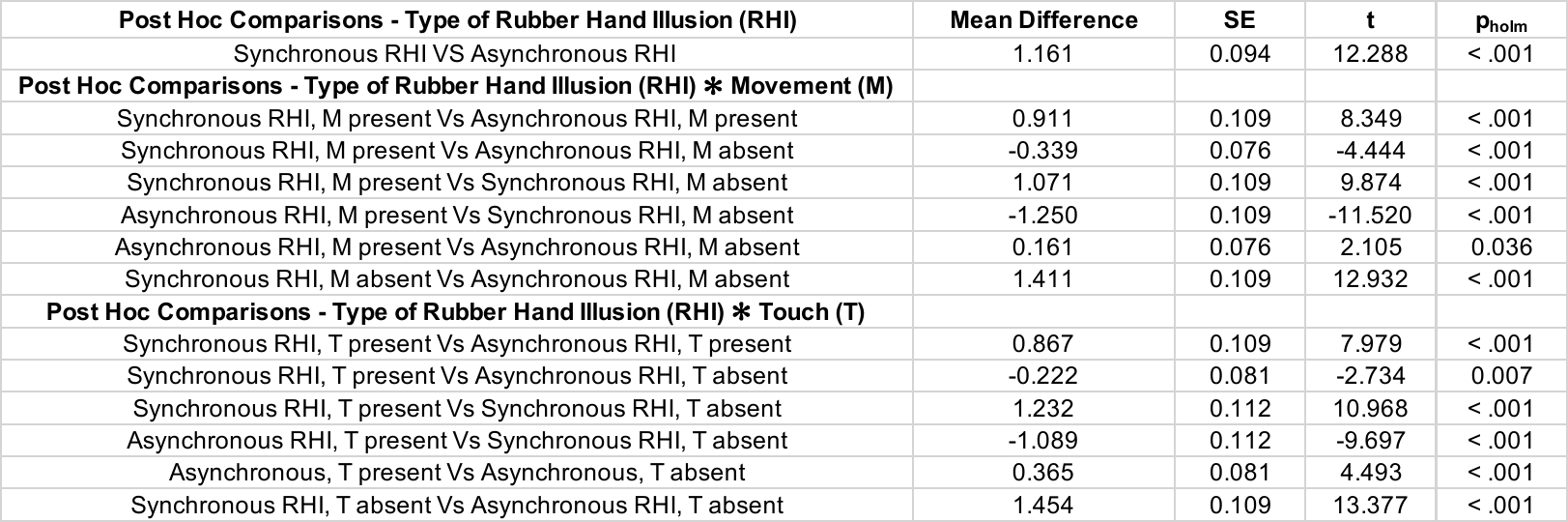

**Supplemental Table 9:** Custom contrast of Experiment 3

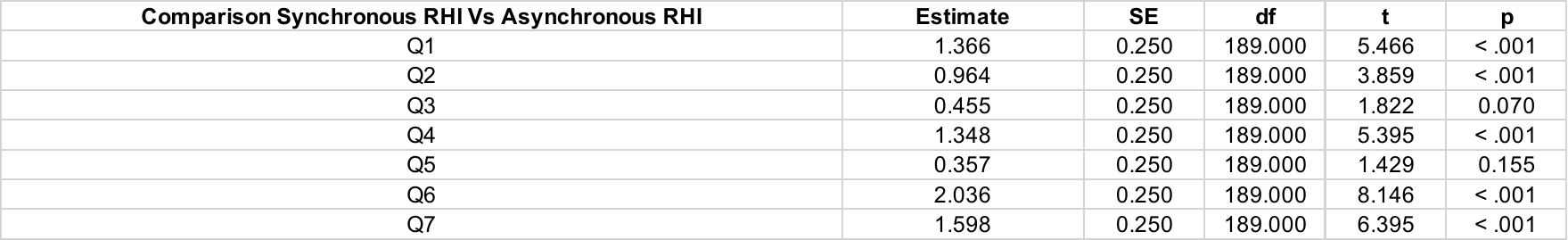

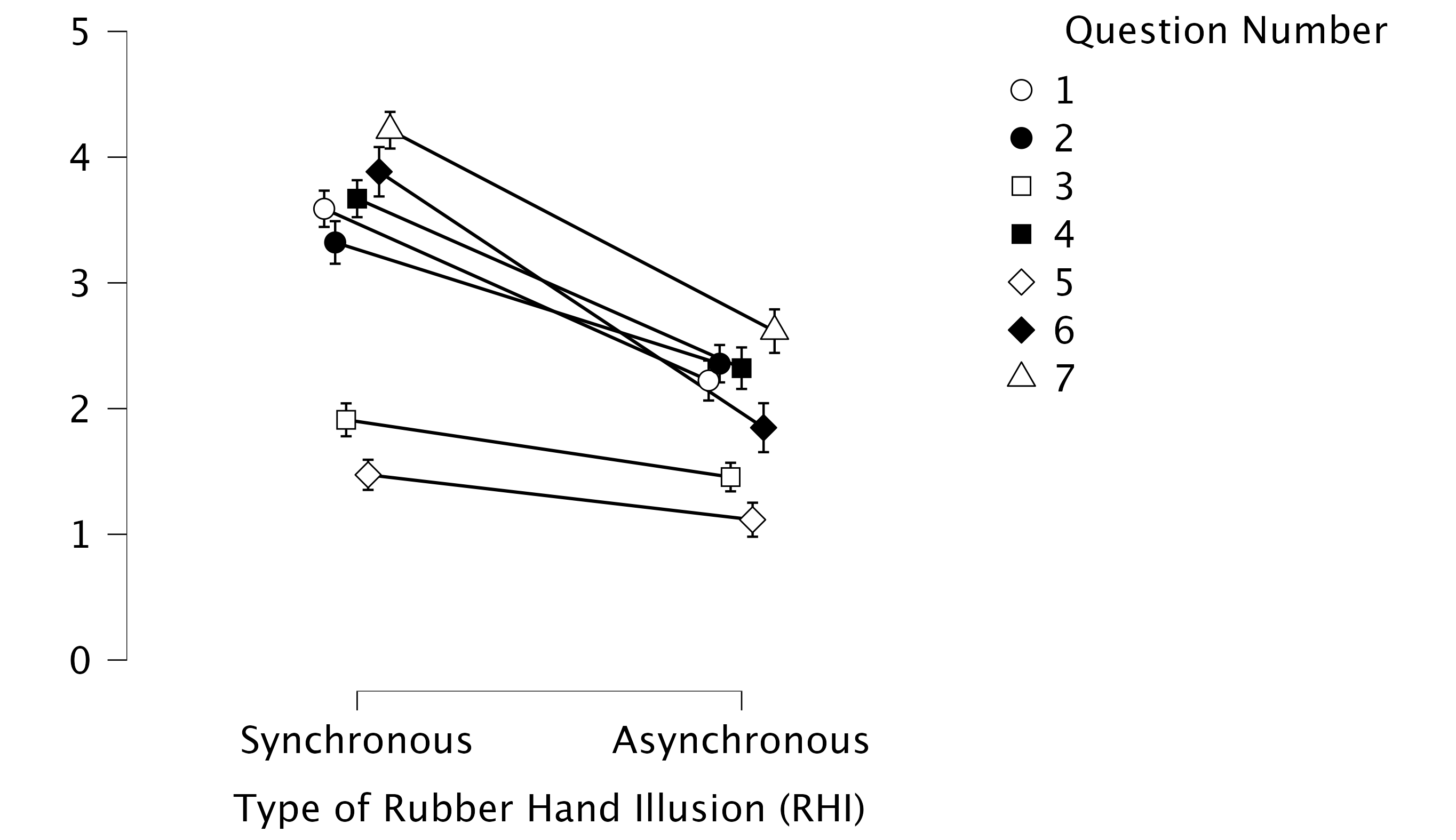

**Supplemental Figure 3:** The plot shows the mean score and the standard error of each question scores in different RHI conditions of Experiment 3.

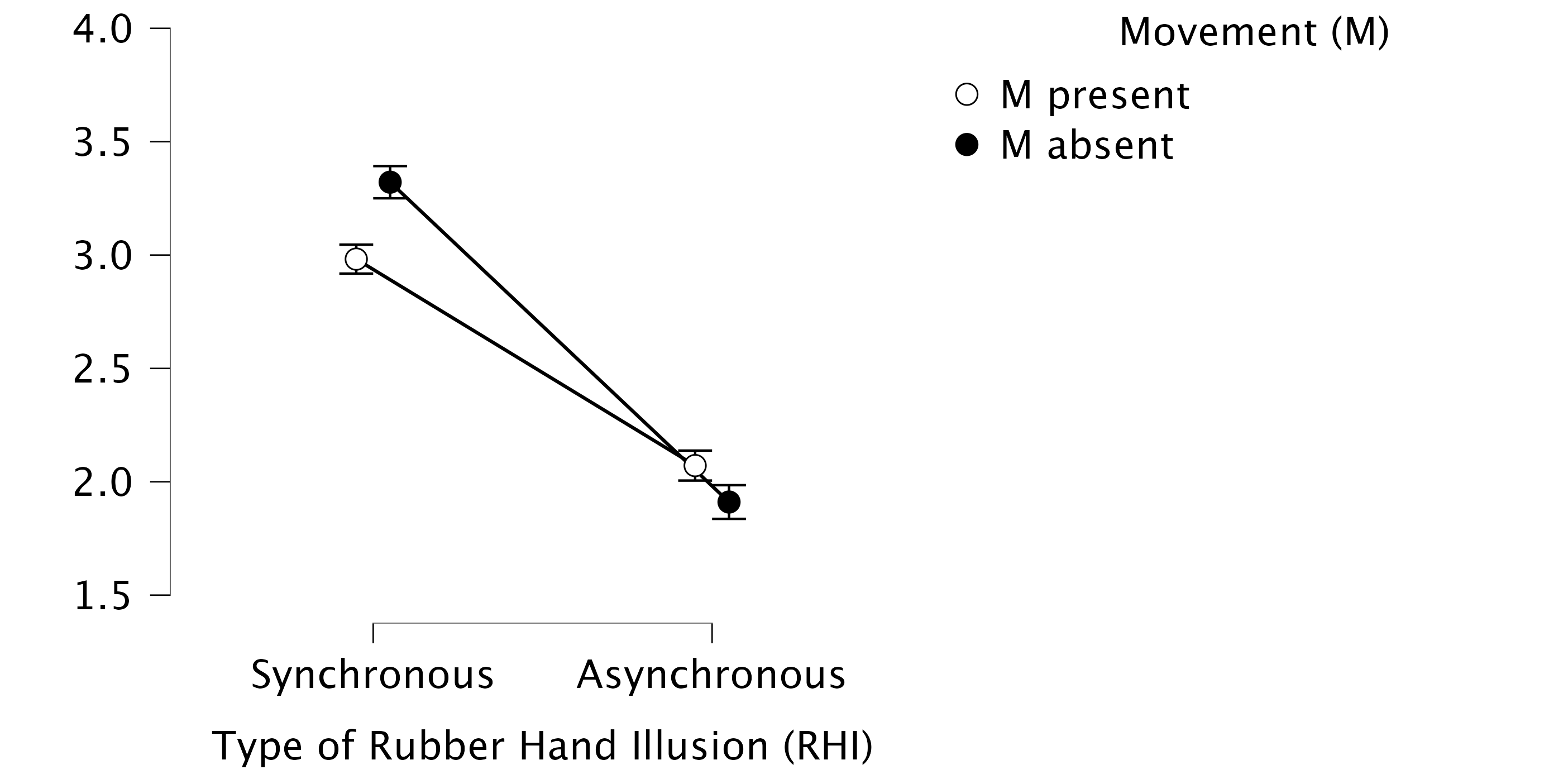

**Supplemental Figure 4:** The plot shows the mean score and the standard error of all questions in synchronous and asynchronous RHI conditions in presence or absent of movement during Experiment 3.

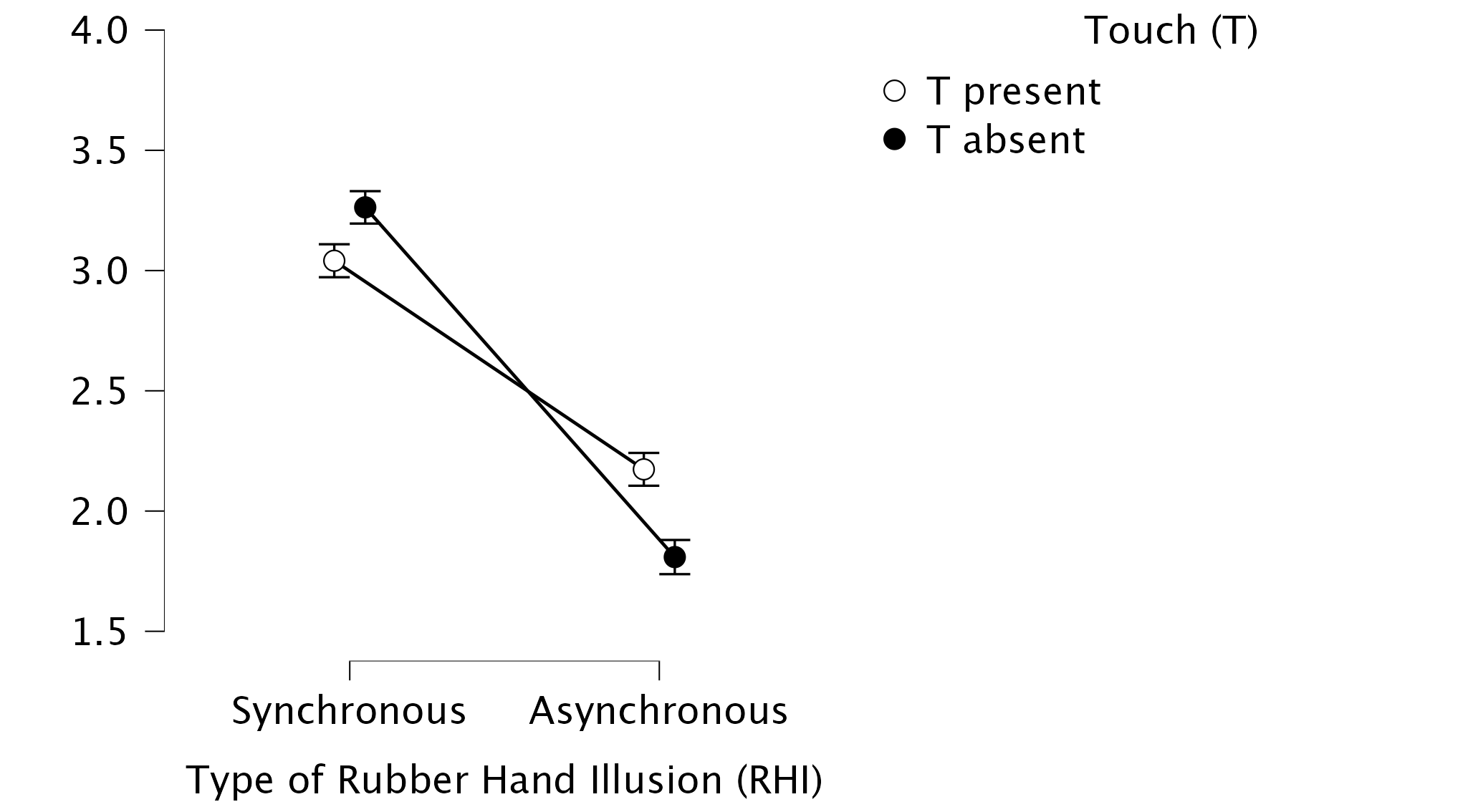

**Supplemental Figure 5:** The plot shows the mean score and the standard error of all questions in synchronous and asynchronous RHI conditions in presence or absent of touch during Experiment 3
